# Science Game Lab: tool for the unification of biomedical games with a purpose

**DOI:** 10.1101/156141

**Authors:** Benjamin M. Good, Sarah Santini, Margaret Wallace, Nicholas Fortugno, John Szeder, Patrick Mooney, Jerome Waldispuhl, Ginger Tsueng, Andrew I Su

## Abstract

Games with a purpose and other kinds of citizen science initiatives demonstrate great potential for advancing biomedical science and improving STEM education. Articles documenting the success of projects such as Fold.it and Eyewire in high impact journals have raised wide interest in new applications of the distributed human intelligence that these systems have tapped into. However, the path from a good idea to a successful citizen science game remains highly challenging. Apart from the scientific difficulties of identifying suitable problems and appropriate human-powered solutions, the games still need to be created, need to be fun, and need to reach a large audience that remain engaged for the long-term. Here, we describe Science Game Lab (SGL) (https://sciencegamelab.org), a platform for bootstrapping the production, facilitating the publication, and boosting both the fun and the value of the user experience for scientific games with a purpose.

## Introduction

Ever since the Fold.it project famously demonstrated that teams of human game players could often outperform supercomputers at the challenging problem of 3d protein structure prediction, so-called ‘games with a purpose’ (von Ahn, 2006) have seen increasing attention from the biomedical research community (Good and Su, 2011). A few other games in this genre include: Phylo for multiple sequence alignment (Kawrykow et al., 2012), EteRNA for RNA structure design (Lee et al., 2014), Eyewire for mapping neural connectivity (Kim et al., 2014), The Cure for breast cancer prognosis prediction (Good et al., 2014), Dizeez for gene annotation (Loguercio, Good and Su, 2013), and MalariaSpot for image analysis (Luengo-Oroz, Arranz and Frean, 2012). Apart from tapping into human intelligence at scale, these efforts have also produced valuable educational opportunities. Many of these games are now used to introduce their underlying concepts in classroom settings (Farley, 2013) where games in all forms are increasingly working their way into curriculums. Concomitant with the rise of these ‘serious games’, citizen science efforts such as the Zooniverse (Segal et al., 2016) and Mark2Cure (Tsueng et al., 2016) have sought similar aims but have packaged their work as volunteer tasks, analogous to unpaid crowdsourcing tasks, rather than as elements of games.

Many of these initiatives have succeeded in independently addressing challenging technical problems through human computation [von ahn], improving science education, and generally raising scientific awareness. However, with so much interest from the scientific community and a booming ecosystem of game developers (Superdataresearch.com, 2017), there are actually relatively few of these games in operation now. Recognizing the opportunity, various groups have attempted to push the area forward through new funding opportunities (e.g. (Grants.nih.gov, 2017)) and through various ‘game jams’ such as the one that produced the game ‘genes in space’ for use in analyzing microarray data in cancer (Coburn, 2014). Here, we take a different approach towards expanding the ecosystem of games with a scientific purpose. Rather than attempting to seed the genesis of specific new game-changing games, we hope to lower the barrier to entry for new games and related citizen science tasks to generally promote the development of the entire field. With this high-level aim in mind, we developed Science Game Lab (SGL) to make it easier for developers to create successful scientific games or game-like learning and volunteer experiences. Specifically, SGL is intended to address the challenges of recruiting players and volunteers, keeping them engaged for the long term, and reducing the development costs associated with creating a scientific gaming experience.

## The Science Game Lab Web application

SGL is a unique, open-source portal supporting the integration of games and volunteer experiences meant to advance science and science education (https://sciencegamelab.org). Unlike other related sites that act more like volunteer management and/or project directory services, such as SciStarter and Science Game Center, SGL is not simply a listing of related websites. Rather, it is an attempt to create a user experience that takes place directly within the SGL context yet still incorporates content from third parties. The system is largely inspired by game industry portals such as Kongregate that enable developers to incorporate their games directly into a unified metagame experience (Patel, 2006).

**Players** can use the portal to find and play games with their achievements within the games tracked on site-wide high score lists and achievement boards (Figure 1). Players can earn the SGL points that drive these leaderboards for actions taken in different games. In this way, SGL provides developers with access to a metagame that can be used to encourage players in addition to the incentives offered within individual games (Figure 2). This metagame can also be used by the system administrators to help direct the player community’s attention to particular games or particular tasks within games. For example, actions taken on new games might earn more points than actions taken on more established games as a way to ‘spread the wealth’ generated by successful games.

**Figure 1.**
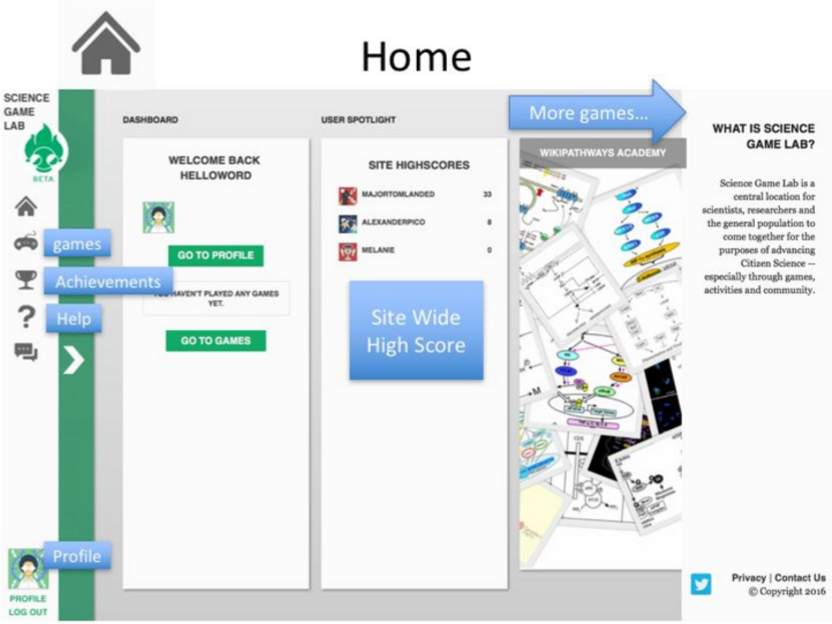
SGL home page demonstrating site-wide high score list, game listing, and links to achievements, help, and user profile information.

**Figure 2.**
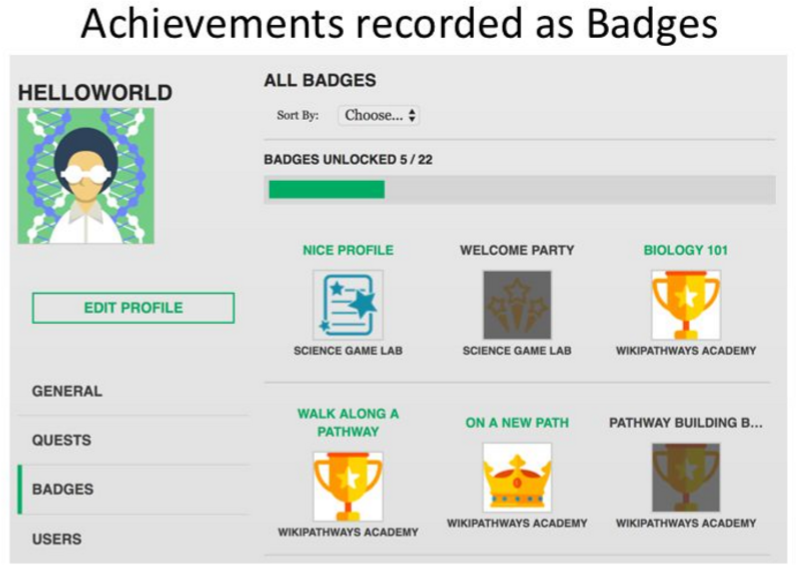
Badges displayed on user’s profile page. Available badges not yet achieved are greyed out.

**Developers** interact with SGL by incorporating a small javascript library into their application and using the SGL ‘developer dashboard’ to pair up events in their game with points, badges and quests managed by the SGL server (Mooney, 2016). At this time, SGL only supports games that operate online as Web applications. The games are hosted by the developers and rendered in the SGL context within an iframe (Raggett, Le Hors and Jacobs, 1999). The SGL iframe provides a ‘heads up display’ that provides real time feedback to game players with respect to events sent back to the SGL server such as earning points, gathering badges, or progressing through the stages of a quest (Figure 3). This display provides developers with the ability to add game mechanics to sites that are not overtly games. For example, Wikipathways incorporated a pathway editing tutorial into SGL, using the heads up display to reward users with SGL points and badges for completing various stages of the tutorial. The tutorial also took advantage of the SGL quest-building tool (Figure 4). Games are submitted by developers for approval by SGL administrators. Once approved, the games appear in the public view and can be accessed by any player.

**Figure 3.**
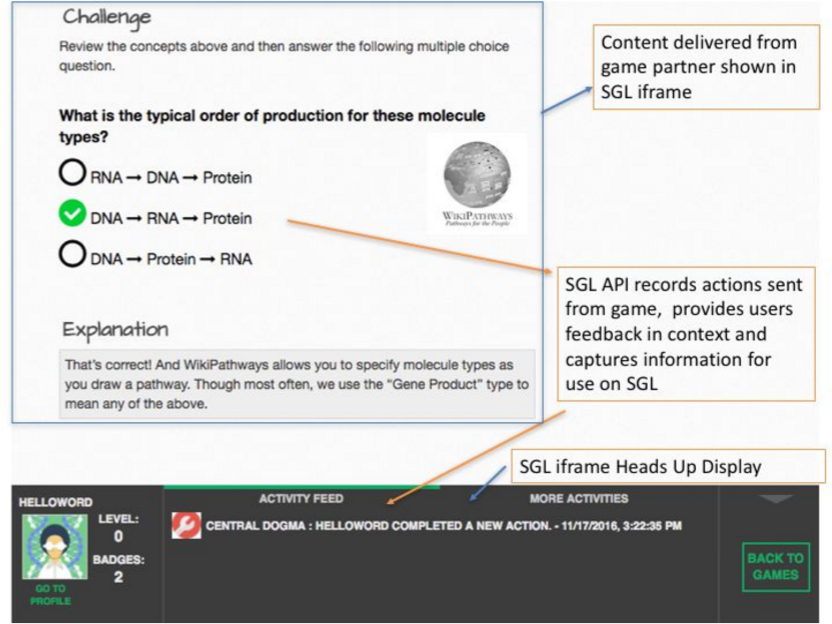
The heads up display provided by the SGL iframe. Shows events captured by the API and provides users with immediate feedback.

**Figure 4.**
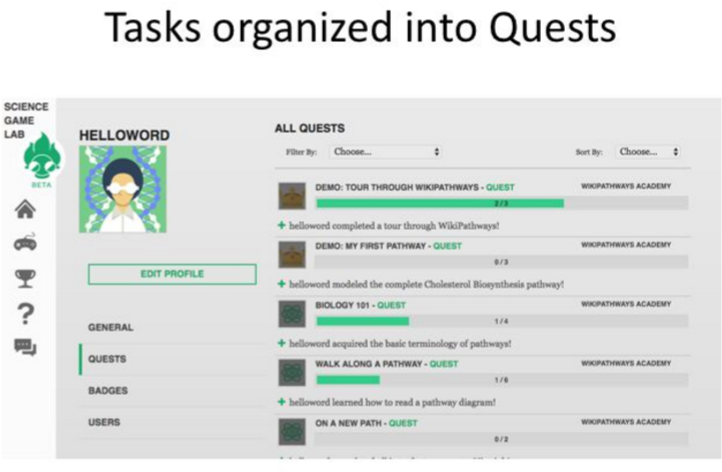
Tasks in SGL can be grouped into quests. The figure shows a particular user’s progress through various quests available within the system.

## Discussion

If a critical initial mass of effective games can be integrated, SGL could strongly benefit new developers by providing immediate access to a large player population. Site-level status, identity and community features can help with the even greater challenge of long-term player engagement, a noted problem in the field (Rallapalli et al., 2015). Within the context of science-related gaming, such status icons might eventually be used as practically useful, real-world marks of achievement inline with the notion of ‘Open Badges’ (Jovanovic and Devedzic, 2014). As demonstrated by the Wikipathways tutorial application, SGL can be used to replace the need for developers to host their own login systems, user tracking databases, and reward systems – all of which can be accomplished using the SGL developer tools. Citizen scientists are not homogenous in their motivations (Lieberoth, 2014). Designing to be inclusive of gamers and non-gamers can be challenging (Bowser, Anne. “Gamifying Citizen Science: Lessons and Future Directions.” (2013).) By offering an alternative means of experiencing a web-based citizen science application, SGL allows developers to cater to both their gaming and non-gaming contributor audience.Together, these features unite to raise the overall potential for growth within the world of citizen science and scientific gaming.

## Future directions

SGL is currently functional, but so far has attracted only a small number of developers willing to integrate their content into the portal. Future work would need to address the challenge of raising the perceived value of integration with the site while lowering the perceived difficulty. Looking forward, key challenges for the future of SGL include better support for:

- games meant for mobile devices
- development of quests that span multiple games
- teachers to build SGL-focused lesson plans and track student progress
- creating new ‘SGL-native’ games
- integration with external authentication systems

None of these are insurmountable challenges, but they all require significant continued investment in software development. As an open source project (https://bitbucket.org/account/user/sulab/projects/SGL), we encourage contributions from anyone that shares in our vision of spreading and doing science through the grand unifying principle of fun.

## References

1. Coburn, C. (2014). Play to Cure: Genes in Space. The Lancet Oncology, 15(7), p.688. doi:10.1016/s1470-2045(14)70259-1 http://www.thelancet.com/journals/lanonc/article/PIIS1470-2045%2814%2970259-1/fullte xt?rss=yes

2. Farley, P. (2013). Using the Computer Game “FoldIt” to Entice Students to Explore External Representations of Protein Structure in a Biochemistry Course for Nonmajors. Biochemistry and Molecular Biology Education, 41(1), pp.56–57. doi:10.1002/bmb.20655 http://onlinelibrary.wiley.com/doi/10.1002/bmb.20655/full

3. Good, B. and Su, A. (2011). Games with a scientific purpose. Genome Biology, 12(12), p.135. doi:10.1186/gb-2011-12-12-135 https://genomebiology.biomedcentral.com/articles/10.1186/gb-2011-12-12-135

4. Good, B., Loguercio, S., Griffith, O., Nanis, M., Wu, C. and Su, A. (2014). The Cure: Design and Evaluation of a Crowdsourcing Game for Gene Selection for Breast Cancer Survival Prediction. JMIR Serious Games, 2(2), p.e7. doi:10.2196/games.3350 http://games.jmir.org/2014/2/e7/

5. Grants.nih.gov. (2017). RFA-CA-15-006: Big Data to Knowledge (BD2K) Advancing Biomedical Science Using Crowdsourcing and Interactive Digital Media (UH2). [online] Available at: https://grants.nih.gov/grants/guide/rfa-files/RFA-CA-15-006.html [Accessed 26 Jun. 2017 ].

6. Jovanovic, J. and Devedzic, V. (2014). Open Badges: Novel Means to Motivate, Scaffold and Recognize Learning. Technology, Knowledge and Learning, 20(1), pp.115–122. doi:10.1007/s10758-014-9232-6 http://link.springer.com/article/10.1007/s10758-014-9232-6

7. Kawrykow, A., Roumanis, G., Kam, A., Kwak, D., Leung, C., Wu, C., Zarour, E., Sarmenta, L., Blanchette, M. and Waldispühl, J. (2012). Phylo: A Citizen Science Approach for Improving Multiple Sequence Alignment. PLoS ONE, 7(3), p.e31362. doi: 10.1371/journal.pone.0031362 http://journals.plos.org/plosone/article?id=10.1371/journal.pone.0031362

8. Kim, J., Greene, M., Zlateski, A., Lee, K., Richardson, M., Turaga, S., Purcaro, M., Balkam, M., Robinson, A., Behabadi, B., Campos, M., Denk, W. and Seung, H. (2014). Space–time wiring specificity supports direction selectivity in the retina. Nature, 509(7500), pp.331–336. doi:10.1038/nature13240 https://www.nature.com/nature/journal/v509/n7500/full/nature13240.html

9. Lee, J., Kladwang, W., Lee, M., Cantu, D., Azizyan, M., Kim, H., Limpaecher, A., Gaikwad, S., Yoon, S., Treuille, A. and Das, R. (2014). RNA design rules from a massive open laboratory. Proceedings of the National Academy of Sciences, 111(6), pp.2122–2127. doi: 10.1073/pnas.1313039111 http://www.pnas.org/content/111/6/2122

10. Lieberoth, A. (2014). Getting humans to do quantum optimization – user acquisition, engagement and early results from the citizen cyberscience game Quantum Moves. Human Computation, 1(2). doi:10.15346/hc.v1i2.11 https://arxiv.org/ftp/arxiv/papers/1506/1506.08761.pdf

11. Loguercio, S., Good, B. and Su, A. (2013). Dizeez: An Online Game for Human Gene-Disease Annotation. PLoS ONE, 8(8), p.e71171. doi:10.1371/journal.pone.0071171 http://journals.plos.org/plosone/article?id=10.1371/journal.pone.0071171

12. Luengo-Oroz, M., Arranz, A. and Frean, J. (2012). Crowdsourcing Malaria Parasite Quantification: An Online Game for Analyzing Images of Infected Thick Blood Smears. Journal of Medical Internet Research, 14(6), p.e167. doi:10.2196/jmir.2338 http://www.jmir.org/2012/6/e167/?trendmd-shared=0

13. Mooney, P. (2016). SGL Support – SGL Support – Confluence. [online] Sciencegamelab.atlassian.net. Available at: https://sciencegamelab.atlassian.net/wiki/display/SGL1/SGL+Support [Accessed 26 Jan. 2017].

14. Patel, R. (2006). Kongregate: a Next Generation Web Games Marketplace. [online] TechCrunch. Available at: https://techcrunch.com/2006/10/19/kongregate-a-next-generation-web-games-marketplace/ [Accessed 26 Jan. 2017].

15. Raggett, D., Le Hors, A. and Jacobs, I. (1999). Frames in HTML documents. [online] W3.org. Available at: https://www.w3.org/TR/1999/REC-html401-19991224/present/frames.html#h-16.5 [Accessed 26 Jan. 2017].

16. Rallapalli, G., Saunders, D., Yoshida, K., Edwards, A., Lugo, C., Collin, S., Clavijo, B., Corpas, M., Swarbreck, D., Clark, M., Downie, J., Kamoun, S. and MacLean, D. (2015). Lessons from Fraxinus, a crowd-sourced citizen science game in genomics. eLife, 4. doi:10.7554/eLife.07460 https://elifesciences.org/content/4/e07460

17. Segal, A., Gal, Y., Kamar, E., Horvitz, E., Bowyer, A. and Miller, G. (2016). Intervention strategies for increasing engagement in crowdsourcing: platform, predictions, and experiments. In: G. Brewka, ed., IJCAI’16 Proceedings of the Twenty-Fifth International Joint Conference on Artificial Intelligence. [online] New York, New York: AAAI Press, pp.3861–3867. Available at: https://www.ijcai.org/Proceedings/16/Papers/543.9df [Accessed 26 Jan. 2017 ].

18. Superdataresearch.com. (2017). SuperData Research | Games data and market research » Market Brief — Year in Review 2016. [online] Available at: https://www.superdataresearch.com/market-data/market-brief-year-in-review/ [Accessed 26 Jan. 2017 ].

19. Tsueng, G., Nanis, S., Fouquier, J., Good, B. and Su, A. (2016). Citizen Science for Mining the Biomedical Literature. Citizen Science: Theory and Practice, 1(2), p.14. doi:10.5334/cstp.56 http://theoryandpractice.citizenscienceassociation.org/articles/10.5334/cstp.56/

20. von Ahn, L. (2006). Games with a Purpose. Computer, 39(6), pp.92–94. doi:10.1109/MC.2006.196 http://ieeexplore.ieee.org/document/1642623/

